## Supplemental Table 1 and Figure 1 for "Torque Teno Sus Virus 1: A Potential Surrogate Pathogen to Study Pig-Transmitted Transboundary Animal Diseases"

Table S3. Comparison of three molecular clock models used in BEAST.

|  | Molecular clock model | Particle # in nested sampling | Marginal likelihood | Standard deviation |
| --- | --- | --- | --- | --- |
| 1 | Strick clock | 30 | -2973.39 | 1.97 |
| 2 | Random local clock | 30 | -2973.44 | 1.97 |
| 3 | Optimized relaxed clock | 30 | -2971.08 | 1.93 |

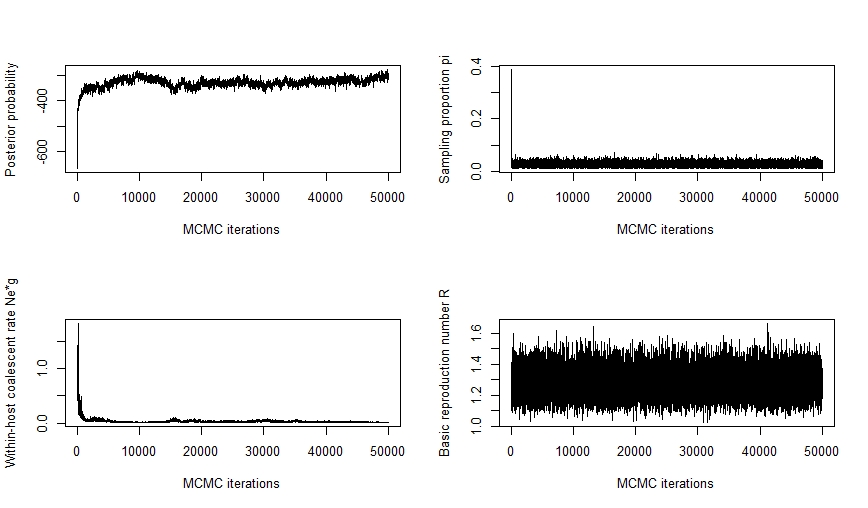

Figure S1. Trace plot of model parameters through MCMC iterations used in TransPhylo.
